# Deletion of *Trpm4* alters the function and expression of Na_V_1.5 channel in murine cardiac myocytes

**DOI:** 10.1101/2020.07.06.188961

**Authors:** Ozhathil Lijo Cherian, Jean-Sébastien Rougier, Prakash Arullampalam, Maria C. Essers, Hugues Abriel

**Affiliations:** Institute of Biochemistry and Molecular Medicine, and Swiss National Centre of Competence in Research (NCCR) TransCure, University of Bern, Bühlstrasse 28, 3012, Bern, Switzerland

## Abstract

Transient receptor potential melastatin member 4 (TRPM4) encodes a Ca^+^ -activated non-selective cation channel that is functionally expressed in several tissues including the heart. Pathogenic mutants in *TRPM4* have been reported in patients with inherited cardiac diseases including conduction block and Brugada syndrome. Heterologous expression of mutant channels in cell lines indicates that these mutations can lead to an increase or decrease in TRPM4 expression and function at the cell surface. While the expression and clinical variant studies further stress the importance of TRPM4 in cardiac function, the cardiac electrophysiological phenotypes in *Trpm4* knockdown mouse models remain incompletely characterized. To study the functional consequences of *Trpm4* deletion on cardiac electrical activity in mice, we performed perforated-patch clamp and immunohistochemistry studies on isolated atrial and ventricular cardiac myocytes and surface, pseudo and intracardiac ECGs either *in vivo* or on Langendorff-perfused explanted mouse hearts. We observed that Trpm4 is expressed in atrial and ventricular cardiac myocytes and that deletion of *Trpm4* unexpectedly reduces the peak Na^+^ currents in the myocytes. Hearts from *Trpm4^-/-^* mice presented increased sensitivity towards mexiletine, a Na^+^ channel blocker, and slower intraventricular conduction, consistent with the reduction of peak Na^+^ current observed in the isolated cardiac myocytes. This reduction in Na^+^ current is explained by the observed decrease in protein expression of Na_V_1.5 in *Trpm4^-/-^* mice. This study suggests that Trpm4 expression impacts Na^+^ current in murine cardiac myocytes and points towards a novel function of Trpm4 regulating the Na_V_1.5 expression in murine cardiac myocytes.

## Introduction

The cardiac Ca^2+^-activated non-selective cation (NSca) currents were first measured in cultured rat neonatal myocytes in the early 1980s [1]. The molecular identity of these current components remained largely unknown until a member of TRPM family, TRPM4b, was cloned [2–5], which was found to share the biophysical properties of a native NSca current from human atrial myocytes [6]. TRPM4 belongs to the transient receptor potential (TRP) ion channels superfamily, comprising mostly large Ca^2+^-permeable cation channels that are expressed in different tissue types and are activated by a broad spectrum of physicochemical stimuli under varying cellular conditions. TRPM4 and TRPM5 are the only members of the TRP family that are not permeable to divalent cations such as Ca^2+^ or Mg^2+^. Instead, TRPM4 is activated by an increase in intracellular Ca^2+^, and voltage further modulates its gating, resulting in an outwardly rectifying current [7–9]. This distinctive biophysical feature of TRPM4 could allow Na^+^ entry at negative potentials leading to membrane depolarization, and K^+^ efflux at positive potentials facilitating membrane repolarization.

In human hearts, the expression of TRPM4 has been detected in Purkinje fibers, atria, and ventricles [10]. Clinical relevance of TRPM4 emerged from the identification of the first mutation in patients with hereditary cardiac conduction slowing disorders. This mutation caused the replacement of a glutamate residue at position 7 with lysine (p.E7K), which led to an increase in protein function and surface expression [10]. To date, more than 25 variants of TRPM4 are associated with conduction disorders and Brugada syndrome [11–15]. However, phenotype-genotype correlations with TRPM4 variants remain ambiguous as both gain- and loss-of-function were reported to have overlapping clinical phenotypes. In one of our recent works, we found that some *TRPM4* mutations altered the protein degradation rate, which led to either increased or decreased stability of the protein at the membrane [13]. How the altered degradation rates are related to the clinical phenotypes of the respective mutations is still unclear. In mouse heart, the role of TRPM4 has been studied by deletion of *Trpm4* [16,17] and inhibition of TRPM4-mediated current using 9-phenanthrol [18–20]. In either case, the cardiac action potentials (APs) were shortened suggesting a role of TRPM4 in the repolarization phase. However, discrepancies exist in these studies regarding their baseline cardiac phenotypes and doubts regarding the lack of specificity of 9-phenanthrol [21–23].

Previous studies have proposed the hypothesis that TRPM4 may functionally affect the activity of other cardiac ion channels, such as the main cardiac volt-age-gated Na^+^ channel Na_V_1.5 [12,24]. Since TRPM4 carries an inward current close to resting membrane potential and an outward current at depolarized potentials, increasing or decreasing TRPM4 expression may critically influence the availability of Na_V_1.5 channels.

In the present study, we evaluated whether TRPM4 indeed has any crosstalk with Na_V_1.5 in cardiac cells and tissues. We investigated the consequences of *Trpm4* deletion on cardiac electrical activity in mice. We electrophysiologically characterized individual isolated cardiac myocytes from atria and ventricles and observed a decrease in the functional expression of Na_V_1.5 due to *Trpm4* deletion. We further demonstrated the impact of *Trpm4* deletion with pseudo- and intracardiac ECGs on Langendorff-perfused explanted hearts. This study provides the first evidence of Trpm4 directly impacting Na^+^ current in murine cardiac myocytes.

## Materials and Methods

### 1) *Trpm4^-/-^* C57BL6/N mice

*Trpm4*^-/-^ mice, previously generated as reported elsewhere[25], were a kind gift from Dr. Rudi Vennekens (KU Leuven) and were backcrossed 10 generations on a C57BL/6N background. For all experiments, male *Trpm4*^-/-^ mice and wildtype (WT) littermates aged 12-15 weeks were used. All experiments involving animals were performed according to the Swiss Federal Animal Protection Law and were approved by the Cantonal Veterinary Administration, Bern. This investigation conforms to the Guide for the Care and Use of Laboratory Animals, published by the US National Institutes of Health (NIH publication no. 85-23, revised 1996).

### 2) Isolation of atrial and ventricular myocyte isolation

Mice were anesthetized with 200 mg/kg ketamine and 20 mg/kg xylazine i.p. and sacrificed by cervical dislocation, after which the heart was rapidly excised, cannulated, and mounted on a Langendorff column for retrograde perfusion at 37°C. The heart was rinsed free of blood for 5 mins with nominally Ca^2+^-free solution containing (mmol/L): 135 NaCl, 4 KCl, 1.2 MgCl_2_, 1.2 NaH_2_PO_4_, 10 HEPES, 11 glucose, pH 7.4 (NaOH). Next, the heart was perfused with the same solutions with 50 *μ*M Ca^2+^ and collagenase type II (1 mg/mL, CLS-2, Worthington, NJ, USA) for 15 minutes. Following digestion, the atria and ventricles were separated and transferred to nominal 100 *μ*M Ca^2+^ solution. For atrial myocytes isolation, only the apex of right atrium was excised and minced into small pieces. The cells were dispersed by gentle pipetting with a fire-polished glass pasteur pipette. The ventricle was transferred to above-mentioned buffer with 100 *μ*M Ca^2+^, minced into small pieces to liberate single ventricular myocytes by gentle pipetting, and filtered through a 100 μm nylon mesh. For electrophysiology studies, isolated myocytes were washed 3 times and the calcium concentration was progressively raised to 1 mmol/L within ~30 minutes. Then the cell suspension was placed on a gently rotating shaker at room temperature 22-25°C until use, within 6 hours after isolation.

### 3) Single-cell myocyte electrophysiology

#### Data acquisition and analysis

Patch-clamp experiments were performed on isolated myocytes in voltage- or current-clamp mode using MultiClamp 700B and VE-2 controlled by Clampex 10 via a digidata 1332A or 1440A series, respectively. Data were exported to IGOR PRO (WaveMetrics, Lake Oswego, OR, USA) for analysis. Patch electrodes were pulled from borosilicate glass capillaries (World Precision Instruments, Germany GmbH) and had a resistance between 2 – 5 MΩ when filled with internal solutions. All experiments were conducted at room temperature (22 – 25°C).

#### Action potential measurements

Action potential (AP) recordings on isolated myocytes were performed in perforated patch configuration. Cardiomyocytes were bathed in a solution containing (mmol/L) 140 NaCl, 5.4 KCl, 1.8 CaCl_2_, 1.2 MgCl_2_, 10 HEPES, and 5 glucose, pH 7.4 (NaOH). Intracellular patch electrodes were filled with (mmol/L) 120 KCl, 1.5 CaCl_2_, 5.5 MgCl_2_, 5 Na_2_ATP, 5 K_2_-EGTA, 10 HEPES and amphotericin B (225 μg/mL), pH 7.4 (KOH). APs were elicited at 0.5 Hz with rectangular pulses (5 ms at 125% threshold) in current-clamp mode in repeated sequences. The APs were analyzed offline for resting membrane potential, maximal upstroke velocity (dV/dt)_max_ and 30, 50 and 90% repolarization durations. Data from 30 consecutive APs were averaged for each cell.

#### Voltage clamp recordings

Peak sodium currents (I_Na_) were measured in wholecell configuration using an internal solution (mmol/L) 60 CsCl, 70 Cs-Aspartate, 1 CaCl_2_, 1 MgCl_2_, 10 HEPES, 11 EGTA and 5 Na_2_ATP, pH 7.2 (CsOH). Isolated myocytes were bathed in solution containing (mmol/L) 10 NaCl, 120 N-methyl-D-glutamine (NMDG-Cl), 1.8 CaCl_2_, 1.2 MgCl_2_, 5 CsCl, 10 HEPES, and 5 glucose, pH 7.4 (CsOH) along with 10 μM CoCl_2_, and 10 μM nifedipine to inhibit Ca^2+^ currents. Peak I_Na_ was measured from a holding potential of −110 mV following steps of 5 mV from −130 to +35 mV with a cycle length of 5 s. Current densities (pA/pF) were calculated by dividing the peak current amplitude by cell capacitance. Series resistance and cell membrane capacitance were compensated for 80%. Voltage dependence of activation was determined from the I/V relationship by normalizing peak I_Na_ to driving force and plotting normalized conductance vs. membrane voltage (V_m_). Voltage dependence of steady-state inactivation was obtained by plotting the normalized peak current (25 ms test pulse to −20 mV after a 500 ms conditioning pulse) vs. V_m_. Voltage dependence of activation and inactivation curves were fitted with the Boltzmann function ((V_m_) = 1 / (1 + exp[(V_m_ – V_1/2_) / k])), where V_1/2_ is the half-maximal voltage of (in)activation and k is the slope factor. I_K1_ was measured using the same internal and external solutions as those used for action potential recordings mentioned above without amphotericin B. Na^+^ channels were blocked with 50 μM tetrodotoxin (TTX) and Ca^2+^ channels were blocked with 3 mM cobalt chloride (CoCl_2_) added to the external solution. I_K1_ barium-sensitive current was calculated by subtracting potassium current recorded after perfusion of the extracellular solution containing 100 μM barium chloride (BaCl_2_) to potassium current recorded before application of BaCl_2_.

### 4) Pseudo- and intracardiac ECG recordings on Langendorff-perfused hearts

Mice were anesthetized with 200 mg/kg ketamine and 20 mg/kg xylazine i.p. and sacrificed by cervical dislocation, after which the heart was rapidly excised. Hearts were retrogradely perfused on a Langedorff system using a modified Krebs-Henseleit buffer (KHB) containing (mmol/L) 116.5 NaCl_2_, 25 NaHCO_3_, 4.7 KCl, 1.2 KH_2_PO_4_, 1.2 MgSO_4_, 11.1 glucose, 1.5 CaCl_2_, 2 Na-pyruvate and bubbled with 95% O_2_ and 5% CO_2_ at 37°C. The perfusion pressure was maintained at 70 mmHg. Pseudo-ECGs were recorded using a pair of thin silver electrodes, with the negative end at the root of aorta and the positive on the apex of the ventricles. The data were acquired using Powerlab bioamplifiers and were low-pass filtered at 5 kHz and high-pass filtered at 10 kHz and sampled at 1 k/sec. For the initial 10 mins, the heart was perfused with KHB buffer followed by 40 μg/mL mexiletine or vehicle control (methanol) dissolved in KHB. Pseudo-ECGs were continuously monitored throughout the experiment and later analyzed using LabChart Pro V.8 (AD Instruments, Australia). The shape of the P wave, QRS complex, and QT region were defined as previously reported[26]. To quantify the ECG perturbation by mexiletine, we defined the QRS duration 2 minutes before the start of the mexiletine perfusion as pre-mexiletine and 20 minutes after the start of mexiletine perfusion as post-mexiletine. We also studied atrioventricular (AV) and intra-ventricular (IV) conduction delays using 1.1F (EPR-800, Millar Instruments, Houston, TX) octapolar intracardiac catheter inserted through the right atrium (RA) and advanced into the right ventricle (RV) on the explanted Langendorff-perfused heart. Proper catheter position was verified by visualization of at least six intracardiac electrocardiograms at the level of right atria and ventricle overlaid by the P wave and QRS complex from the pseudo-ECGs on the same explanted heart. A representative ECG trace from a typical intracardiac recording configuration is shown in Figure 5A. To calculate the conduction delay, we measured the time delay between the first derivative negative peaks from different electrodes (CH 1-6) of the catheter. The distance between each electrode on the catheter is 1 mm. AV delay was measured between CH 6 and 5 (1 mm distance), while IV delay was measured between CH 4 and 1 (3 mm distance).

### 5) RNA preparation and real-time quantitative RT-PCR

RNA was extracted from isolated ventricular and atrial myocytes. For atrial myocytes, the left and right atria were separated and pooled from three mice for each side. RNA isolation was performed as reported previously[27]. PCR amplification was performed with TaqMan gene expression assay probes for mouse TRPM4 (Mm01205532-m1), Na_V_1.5 (Mm01342518-m1), and GAPDH(Mm99999915-g1). Relative quantification was performed using the comparative threshold (CT) method (ΔΔCT) after determining the CT values for the reference (GAPDH) and the target genes (TRPM4 and Na_V_1.5) for each sample set.

### 6) Western blot

Proteins from freshly isolated myocytes were prepared by lysing in 0.3 mL ice-cold lysis buffer (50 mM HEPES, 150 mM NaCl, 1 mM EGTA, 1 mM EGTA, 10% glycerol, 1.5 mM MgCl_2_, 10 mM NEM, complete protease inhibitor (Roche diagnostics GmbH ref.11697498001) on a rotating wheel at 4°C overnight. The obtained homogenate was centrifuged at 3000×g at 4°C for 15 mins to remove large cellular fragments. Fifty μL of the supernatant was used for Na_V_1.5 staining with total lysate. The remaining supernatant was ultra-centrifuged at 200,000×g for 45 mins in saccharose buffer (1 mol/L saccharose, 1 mol/L HEPES and complete protease inhibitor). The pellets containing total membrane fractions were solubilized in a cold saccharose buffer. Protein concentrations were determined by Bradford assay using bovine serum albumin (BSA) as a standard. About 80 μg of protein from ventricular myocytes and 25 μg from atrial myocytes were dissolved with 4X LDS sample buffer and 100 mM DTT. The sample was denatured at 37°C for 30 mins and separated on 6% or 9% SDS-PAGE gel for detecting Na_V_1.5 or TRPM4, respectively. Proteins were transferred from the gel to a nitrocellulose membrane (Bio-Rad, USA). The respective proteins were detected with custom rabbit polyclonal antibodies (Na_V_1.5 epitope: DRLPKSDSEDGPRA-LNQLS (Pineda Antibody Service, Berlin, Germany); TRPM4 epitope: VGPEKEQSWIPKIFRKKVC (Pineda, Germany). Calnexin (Sigma-Aldrich., MO, USA., Cat No: C4731) was used to normalize TRPM4 and Na_V_1.5 protein band intensity. All the western blots were quantified using Image Studio Lite software (LI-COR Biosciences, NE, USA).

### 7) Statistical analyses

Data are presented as mean ± SEM. Statistical significance of differences between two means were determined using Student’s t-test if *n* ≥ 5. P values ≤ 0.05 were accepted as significant and represented as # in respective figure panels. The experimenter and person analyzing the data were blindfolded for genotype of the mice. The number of mice and the number of cells used for each experiment are represented as *N* and *n*, respectively, in the figure legends.

## Results

### I. Deletion of *Trpm4* affects the mouse cardiac action potential

Previous studies have reported the expression of Trpm4 in atrial and ventricular tissues from mouse heart [16,17,28]. To evaluate the expression of Trpm4 uniquely in isolated myocytes from different compartments of the heart, we quantified protein mRNA expression using freshly isolated atrial and ventricular myocytes. In line with the reported studies, we observed Trpm4 protein (Figure 1A and B) and mRNA (Figure 1C) expression in atrial and ventricular cells. *Trpm4*^-/-^ mice [28] showed a reduction of the 130 kDa Trpm4 signal both in atria and ventricles. Since a strong expression of Trpm4 has been previously reported in colon, we used it as a positive control in WT. Expectedly, this signal was reduced in *Trpm4*^-/-^ (Figure 1A).

**Figure 1.**
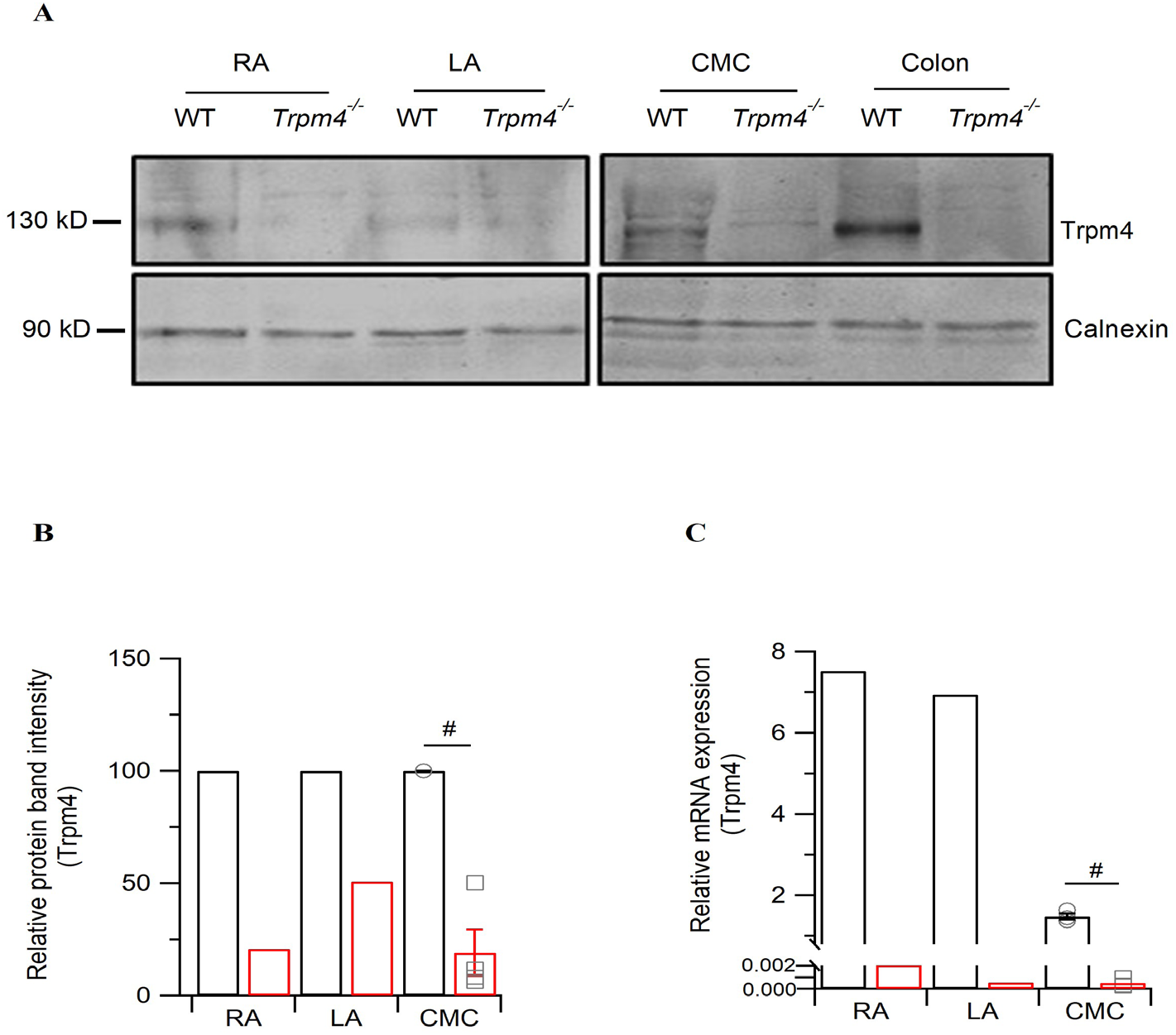
Expression of Trpm4 in mouse atrial and ventricular myocytes. A) Representative immunoblots of Trpm4 protein expression in isolated cardiac myocytes from right (RA) and left (LA) atrium, ventricles (CMC) and colon tissue. B) Quantification of the immunoblot showing protein expression in atrial and ventricular myocytes. *(N* = 4) C) Quantification of *Trpm4* mRNA using RT-qPCR. The Ct values were corrected with GAPDH (Y-axis is split for clarity) (*N* = 3). (For RA and LA, cells were pooled together from all mice for each atrium. For CMC, each mouse was studied individually). (#: *p* < 0.05, WT vs *Trpm4*^-/-^).

To determine the functional consequences of *Trpm4* deletion on cardiac electrical activity at the cellular level, we recorded cardiac action potential (AP) from freshly isolated right atrial and ventricular cardiomyocytes from WT and *Trpm4^-/-^* mice (Figure 2). In AP recordings from atrial cardiomyocytes, we did not observe any difference in upstroke velocity or action potential duration (APD) among the groups (Figure 2A, and 2B). However, the resting membrane potential (RMP) of atrial cardiomyocytes from *Trpm4^-/-^* was significantly depolarized by ~1 mV when compared to the WT (Figure 2B). Since in cardiac myocytes I_K1_ is one of the major components for the RMP, we compared the I_K1_ currents and did not observe any significant alterations in the peak currents (*p* > 0.05, Supplementary 1). We also measured APs in ventricular cardiomyocytes (Figure 2C). Here, we observed a decrease in the upstroke velocity in *Trpm4*^-/-^ mice (*p* = 0.03); whereas neither APDs nor the RMP were altered (Figure 2D).

**Figure 2.**
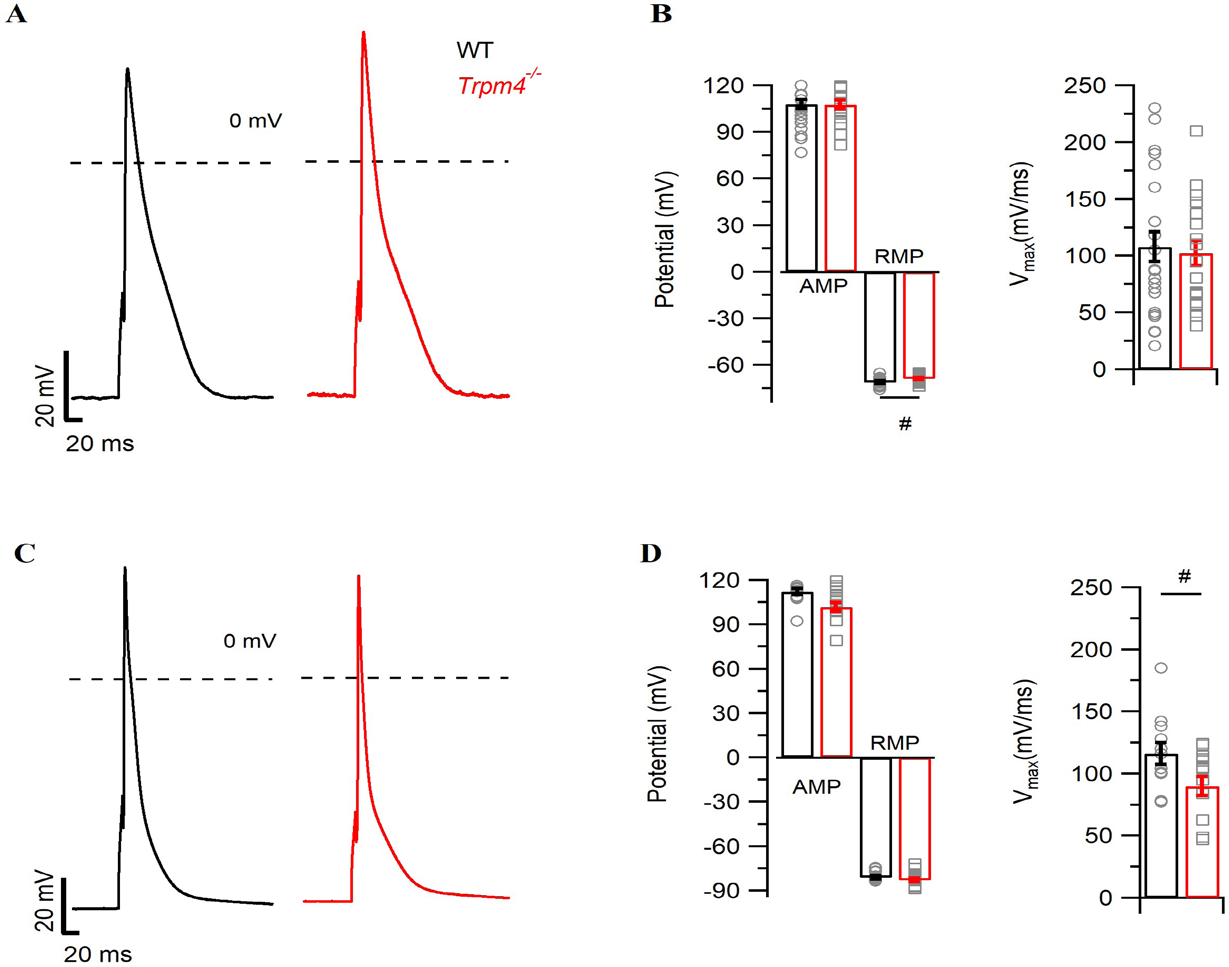
Action potential measurement in iso-lated mouse atrial and ventricular myocytes. Representative traces of AP recorded in WT (black) and *Trpm4^-/-^* (red) mice either from A) right atrial myocytes (*N* = 5, n = 19) or C) ventricular myocytes (*N* = 5, *n* = 14). Average values for APA, RMP, V_max_ and APD at 30, 50 and 90% repolarization (APD_30_, APD_50_ and APD_90_) in WT (black) and *Trpm4^-/-^* (red) either from B) right atrial myocytes or D) ventricular myocytes. (#: *p* < 0.05, WT vs *Trpm4^-/-^*).

### II. Deletion of *Trpm4* affects Na^+^ current in atrial and ventricular myocytes

To study the ionic mechanism contributing to the decrease in the upstroke velocity of the action potential recordings from ventricular myocytes, we measured Na^+^ current using a voltage step protocol in whole-cell configuration. In ventricular cardiomyocytes, the peak Na^+^ current density was reduced by 30% in *Trpm4^-/-^* mice compared to WT (Figure 3A and B, Table 1). Although our AP recordings from atrial myocytes did not reveal any alteration in the upstroke velocity, the peak Na^+^ current in atrial myocytes was reduced by 25% in *Trpm4*^-/-^ compared to WT (Figure 3D and E, Table 1). In addition, the membrane capacitance of atrial myocytes from *Trpm4*^-/-^ was reduced by 20% in both peak Na^+^ current and I_K1_ recordings. (Table 1, Supplementary 1, B). This decrease in the membrane capacitance did not influence the reduction in the peak Na^+^ current observed in atrial myocytes from *Trpm4*^-/-^ mice (Supplementary 2).

**Figure 3.**
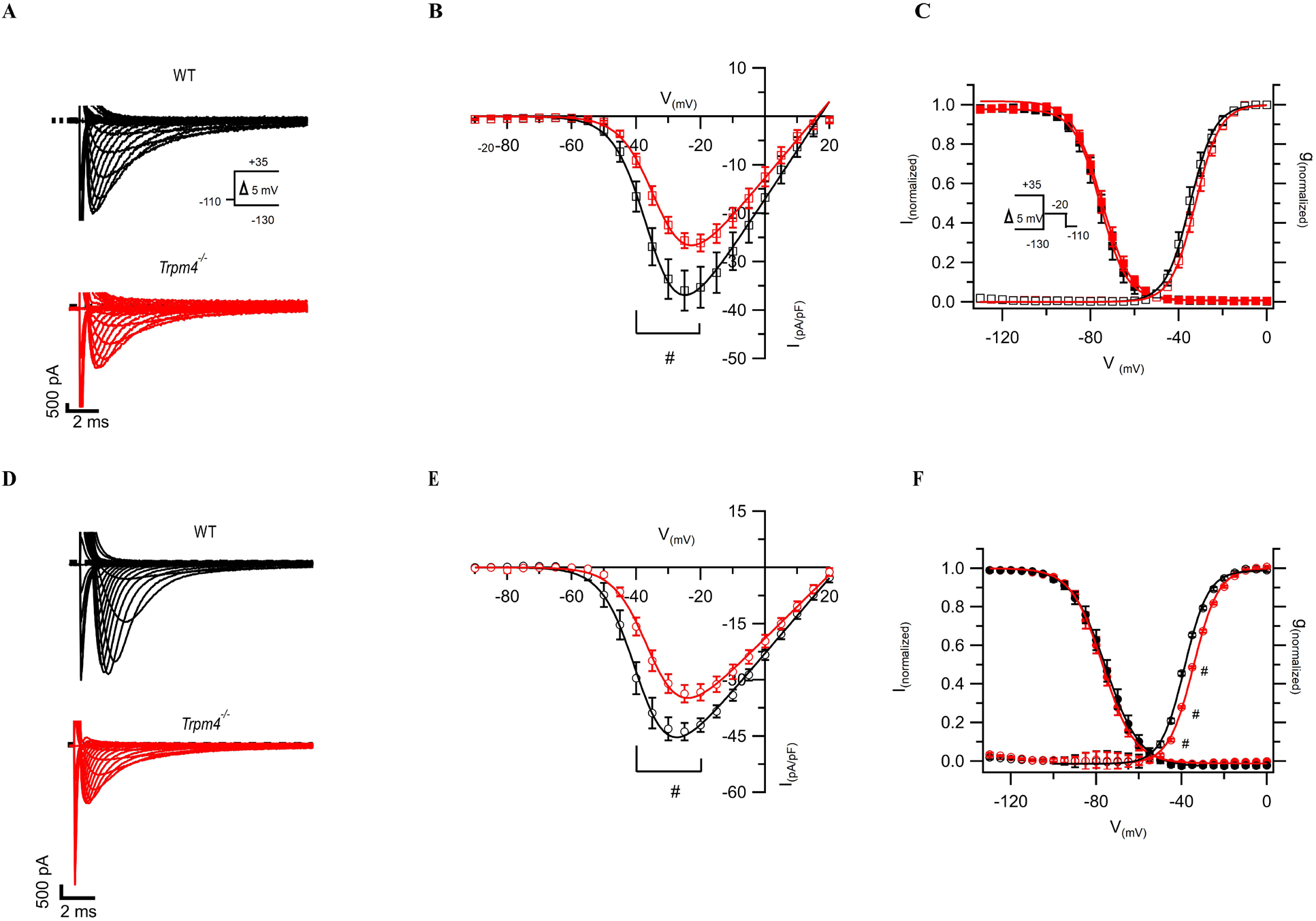
Na^+^ current recordings from isolated cardiomyocytes. A Representative Na^+^ current traces recorded from myocytes in WT (black) or *Trpm4*^-/-^ (red) mice either from ventricular (*N* = 5, *n* = 9 WT, *n* = 12 *Trpm4^-/-^* A) or atrial (*N* = 5, *n* = 20 WT, *n* = 22 Trpm4^-/-^) tissue D) using a voltage step protocol as shown in inset. Na^+^ current-voltage relationship in WT and *Trpm4^-/-^* mice from either ventricular B) or atrial E) cardiomyocytes. C and F show the voltage dependence of activation and inactivation fitted with Boltzmann equation in WT and *Trpm4^-/-^* mice from either ventricular or atrial cardiomyocytes, respectively. (#: *p* < 0.05, WT vs *Trpm4*^-/-^ at a given voltage).

**Table 1.**
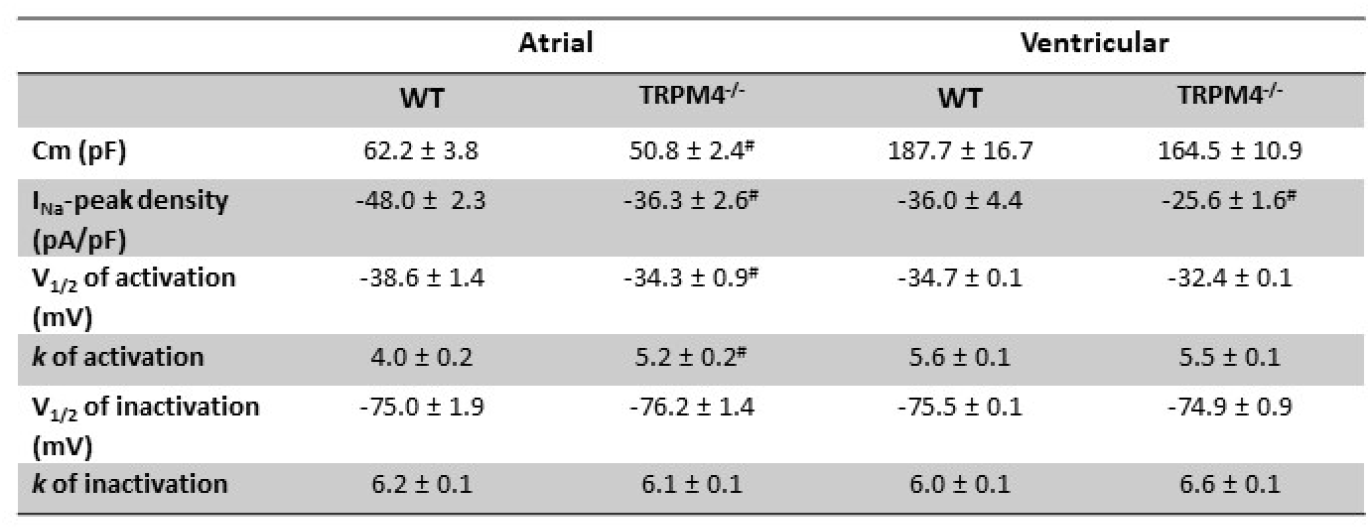
Summary of I_Na_ patch clamp data. (#: *p* ≤ 0.05, WT vs. *Trpm4*^-/-^).

To further assess any alterations in the biophysical properties of the peak Na^+^ current, we measured the V_1/2_ of steady-state activation and inactivation. We observed no differences in V_1/2_ of steady-state inactivation and activation between WT and *Trpm4*^-/-^ ventricular cardiomyocytes (Figure 3 C, Table 1). In atrial myocytes, however, we observed a significant depolarizing shift of 4 mV in the V_1/2_ of steady-state activation in *Trpm4*^-/-^ mice compared to WT, while the steady-state inactivation was not altered (Figure 3 F, Table 1).

Since this reduction of peak Na^+^ current was not previously reported in this mouse line [16], we recently generated in our laboratory a new *Trpm4*^-/-^ mouse strain on a C57BL/6J background (Supplementary 3 and 4) by targeting *Trpm4* exon 10 instead of the exon 15-16 [25]. We measured peak Na^+^ current in WT and *Trpm4*^-/-^ ventricular myocytes and observed a similar decrease of 25% in peak Na^+^ current in these *Trpm4^-/-^* cells (Supple-mentary 5).

### III. *Trpm4*^-/-^ mouse hearts display an increased sensitivity towards the Na^+^ channel inhibitor mexiletine

Since our data indicated an alteration in Na^+^ current density in atrial and ventricular *Trpm4*^-/-^ cardiomyocytes, we next investigated the consequences of Trpm4 deletion on the whole heart by *in vivo* surface ECGs in deeply anesthetized mice. We did not observe any significant alteration in the ECG parameters apart from the heart rate (Table 2). The heart rate in *Trpm4^-/-^* was markedly reduced to 280 ± 19 bpm compared to WT (361 ± 20 bpm; *p* = 0.02).

**Table 2.**
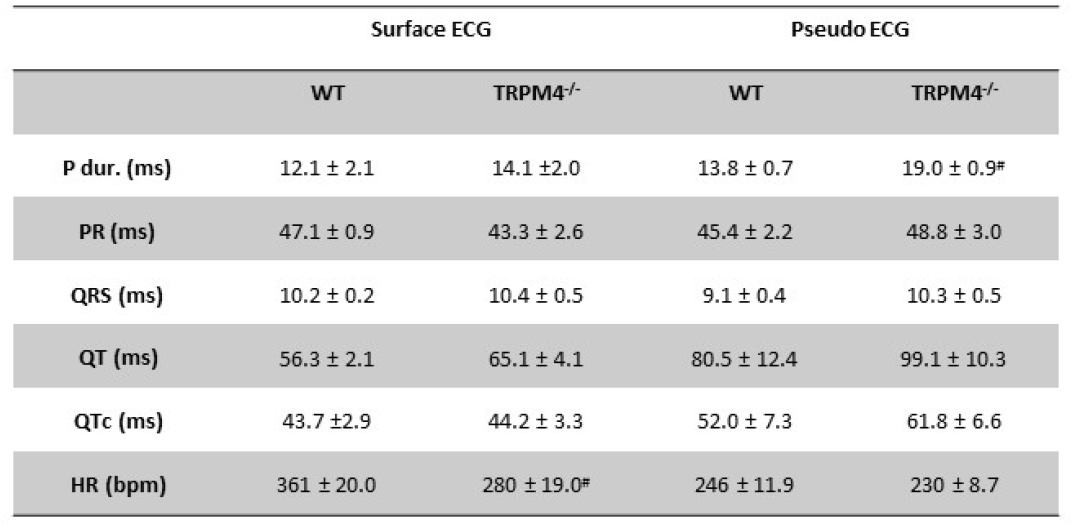
Summary of surface and pseudo-ECG parameters from anesthetized mice.

Further investigations of the electrical properties of *Trpm4*^-/-^ mice were conducted using pseudo ECG recordings on Langendorff-perfused explanted mouse hearts. Such explanted mouse hearts allow perfusion of different drugs while alterations in electrical proper ties can be tracked in real-time. Figure 4A shows an example of a two-lead pseudo-ECG and the analyzed parameters on a perfused explanted WT mouse heart beating spontaneously. In pseudo-ECGs recorded from control buffer perfused hearts, the P duration was significantly increased in *Trpm4^-/-^* compared to WT (*p* = 6*10-^5^) (Figure 4B, Table 2). Other ECG parameters including PR interval, QT, QTc and HR were not altered by *Trpm4* deletion. QRS duration, however, showed a non-significant trend for broadening in *Trpm4^-/-^* hearts (*p* = 0.06) (Figure 4C, Table 2).

**Figure 4.**
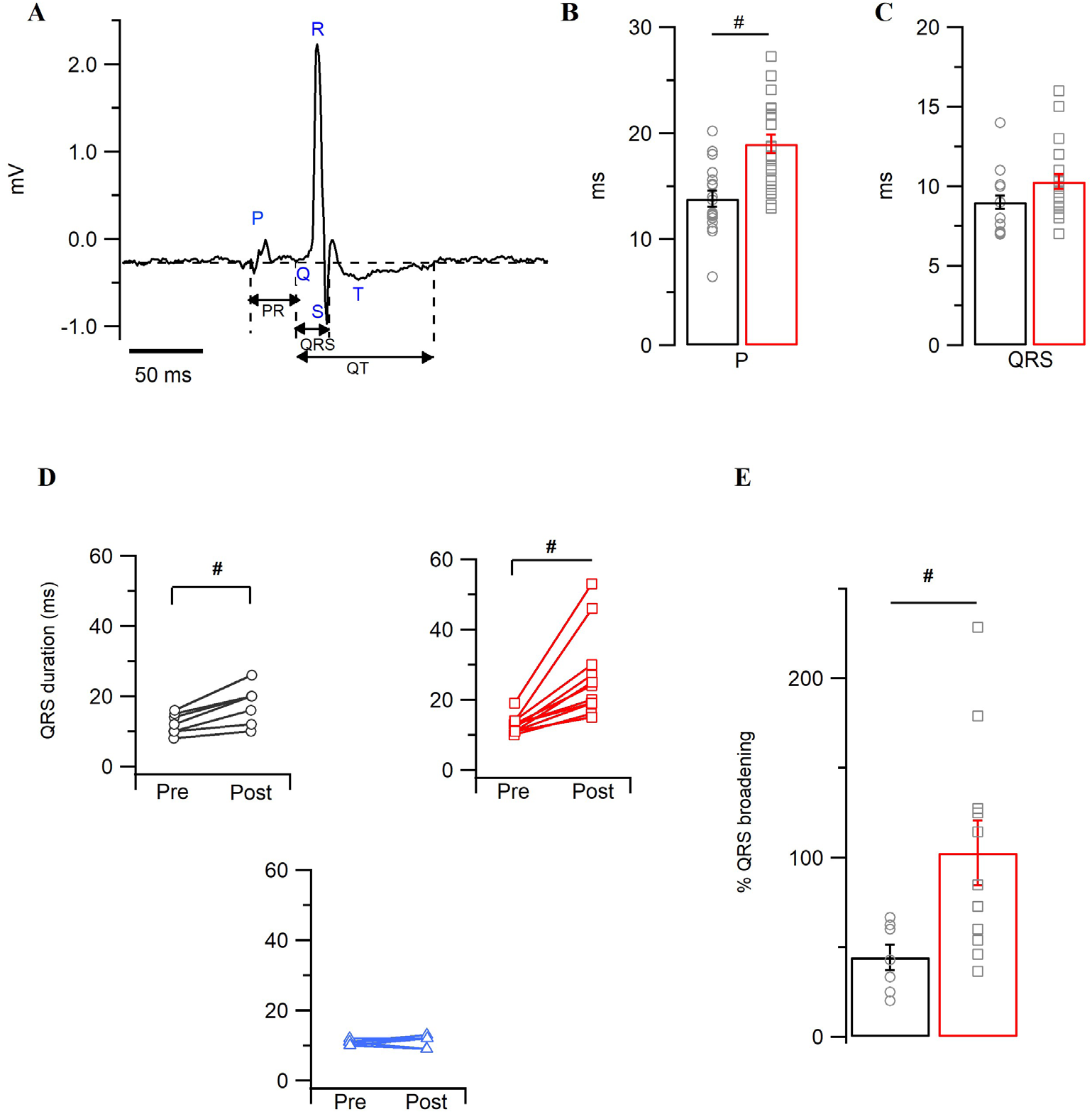
Pseudo-electrocardiogram recording on WT and *Trpm4*^-/-^ explanted hearts. Representative pseudo-ECG trace from WT heart highlighted for different ECG parameters, considered for further comparative studies. Average P (B) and QRS (C) durations recorded from either WT (*N* = 19,black) or *Trpm4*^-/-^ (*N* = 22, red) hearts. D) QRS durations calculated pre- and post-mexiletine perfusion in WT (*N* = 7, black) and *Trpm4*^-/-^ (*N* = 11, red) or perfused with the vehicle control, methanol (*N* = 6, blue). E) The degree of broadening of QRS due to mexiletine perfusion compared between the genotypes as % inhibition. (#: p < 0.05, WT vs. *Trpm4*^-/-^).

To assess whether *Trpm4* deletion affected QRS duration, we challenged the explanted hearts with 40 μg/ mL mexiletine, a Na^+^ channel blocker [29]. Mexiletine broadened the QRS duration in both genotypes (Figure 4D). However, QRS duration from *Trpm4^-/-^* hearts presented a two-fold increase in mexiletine sensitivity compared to WT (QRS broadening (%): 102 ± 18 vs. 44 ± 7, *p* = 0.02) (Figure 4E). The QRS broadening observed in our recordings was exclusively mexile-tine-dependent, as methanol (vehicle control) did not have any effect on the QRS duration (Figure 4D).

### IV. Intra-ventricular conduction is slower in *Trpm4^-/-^* mouse heart than in WT

To investigate any potential conduction delay due to the reduction of peak Na^+^ current in *Trpm4^-/-^* mice, we performed intracardiac ECG measurements on the spontaneously beating explanted hearts using intracardiac catheters with eight electrodes. In most explanted hearts, we could acquire electrical activity from at least six electrodes, one from the right atrium and five from the right ventricle. Pseudo-ECGs were simultaneously acquired, allowing the correct identification of atrial (A) and ventricular (V) peaks in intracardiac ECGs (Figure 5A). The AV delay measured between the A and V signal at an electrode distance of 1 mm did not reveal any difference between the genotypes. However, the intra-ventricular (IV) conduction time from the right ventricle measured at an electrode distance of 3 mm was markedly slower in *Trpm4^-/-^* than in WT hearts (0.95 ± 0.1 vs. 0.3 ± 0.07 ms, *p* = 0.002) (Figure 5C).

**Figure 5:**
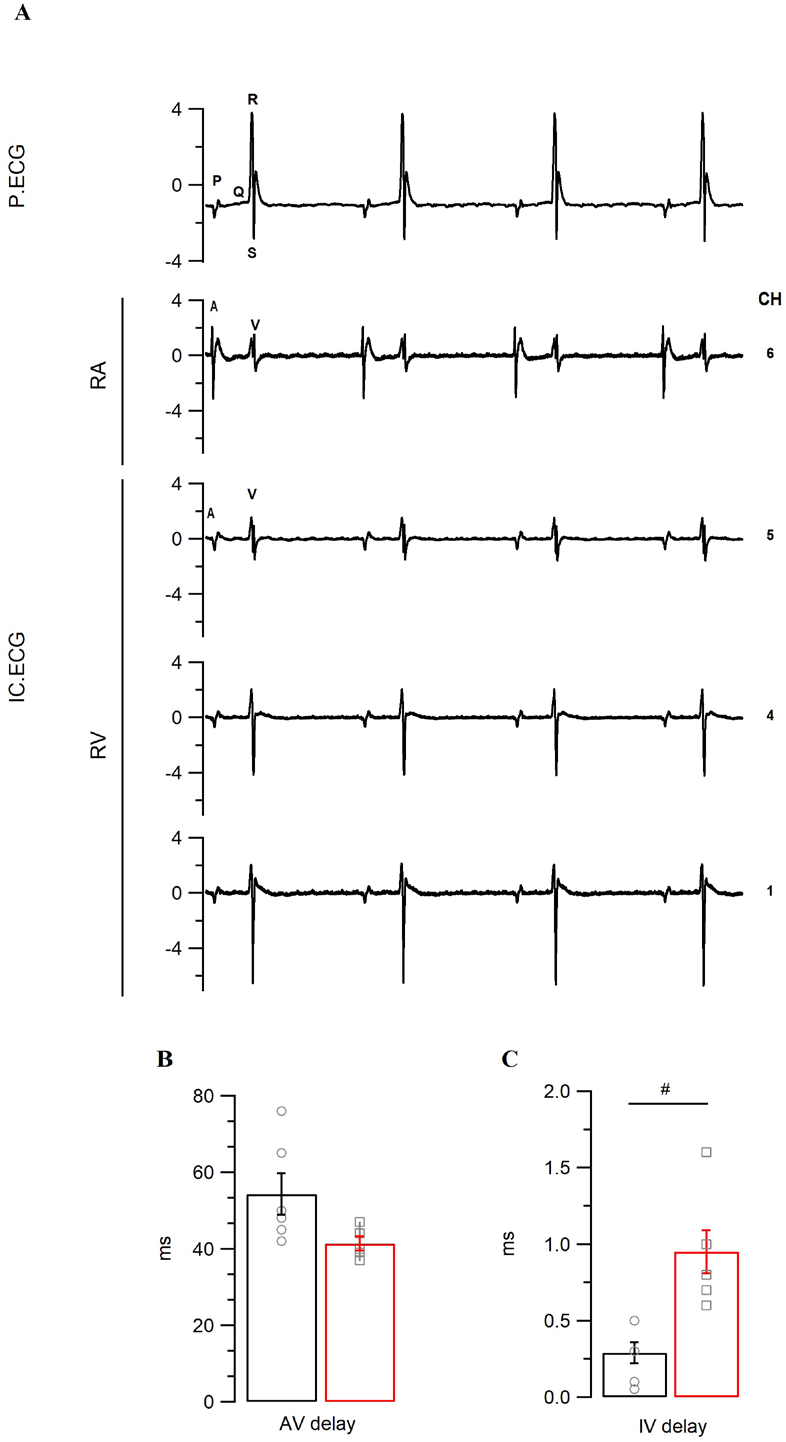
Intracardiac electrocardiogram recordings on WT and *Trpm4*^-/-^ explanted hearts. A) Representative pseudo (P.ECG) and intracardiac (IC.ECG) ECG trace from WT heart. A and V represent respective atrial and ventricle signals from the intracardiac catheter recorded at different electrodes (CH 1-7). Conduction delay either between atria/ ventricle or intraventricular is measured as AV delay (B) or IV delay (C), respectively, and compared between the genotypes. (*N* = 6, #: *p* < 0.05, WT vs. *Trpm4*^-/-^).

### V. Knockdown of *Trpm4* alters Na_V_1.5 expression in mouse ventricles

To further investigate the reduction of peak Na^+^ current observed in isolated atrial and ventricular myocytes, we quantified the expression of Scn5a encoding the cardiac Na^+^ channel Na_V_1.5. mRNA expression levels of Na_V_1.5 in isolated atrial or ventricular cardiac myocytes (Figure 6A) did not differ between WT or *Trpm4*^-/-^. However, the total protein expression of Na_V_1.5 in whole ventricular tissue was significantly reduced by 50% in *Trpm4*^-/-^ compared to WT (Figure 6B and C). In atrial tissue, we observed a slight decrease in the total Na_V_1.5 expression level that we could not statistically validate (Figure 6B and C).

**Figure 6:**
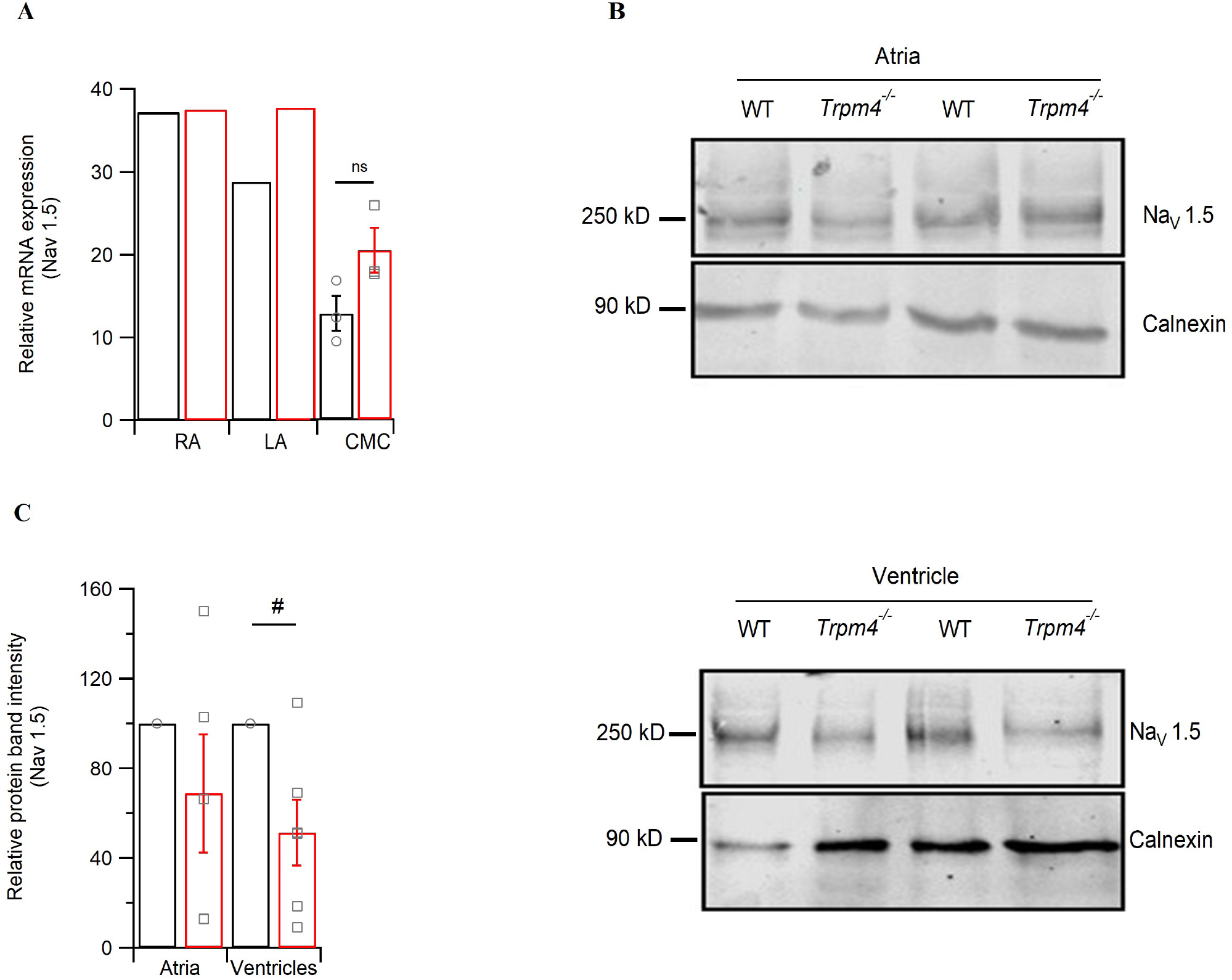
Expression of Na_V_1.5 in mouse atria and ventricles. A) Quantification of Na_V_1.5 mRNA using RT-qPCR on isolated myocytes from RA, LA or CMC. The Ct values were corrected with GAPDH. (*N* = 3; For RA and LA, cells were pooled together from 3 mice. CMCs of 3 mice were studied individually) B) Representative immunoblots of Na_V_1.5 showing total protein expression in either whole atria or ventricles. C) Quantification of the immunoblot showing the total protein expression in whole atria or ventricles (*N* = 6).

## Discussion

In the present study, we compared the functional expression of Trpm4 in freshly isolated murine atrial and ventricular myocytes and studied the consequences of *Trpm4* deletion on mouse heart electrical activity. Here we observed an unexpected decrease in Na_V_1.5 expression and function in *Trpm4*-deficient mouse hearts.

Since the reports in which TRPM4 was identified as the prime molecular candidate for NSca [2,4], functional expression of this channel has been detected in different parts of the heart such as the sino-atrial node, atria, and very low levels in ventricles [6,11,17,20,28,30]. In this study, we quantified the expression of Trpm4 in isolated myocytes from the atria and ventricles. The cardiac myocytes from right atria, and both the ven tricles displayed a stronger Trpm4 protein expression signal; these signals were abolished in *Trpm4*^-/-^ hearts. The functional role of this channel in cardiac electrical activity remains unclear. Trpm4 is suggested to prolong early APDs[16,18], which has been shown using the generic Trpm4 inhibitor 9-phenanthrol [31,32] and in *Trpm4*^-/-^ mice [33–35]. In our perforated-patch AP recordings on isolated myocytes, however, we did not observe any changes in the APDs, while the upstroke velocity in *Trpm4*^-/-^ ventricular myocytes was significantly slower compared to WT. The fast kinetics of Na_V_1.5 underlie the upstroke velocity of a cardiac AP. We, therefore, compared any alteration in the functional expression of Na_V_1.5 due to *Trpm4* deletion in myocytes from atria and ventricles. Indeed, the peak Na^+^ currents conducted by Na_V_1.5 in cardiac myocytes were reduced by 25 and 30% in atria and ventricles, respectively, in *Trpm4*^-/-^ mice compared to WT. The decrease in peak Na^+^ current correlated with a 50% decrease in Na_V_1.5 protein expression in ventricles, while in atria we could not statistically validate any change in Na_V_1.5 expression. However, in atrial myocytes, our study and others reported stronger *Trpm4* expression compared to ventricles [6,17,18,28], the impact of Trpm4 deletion is more complex. In addition to a decrease in the peak Na^+^ current, biophysical properties of the Na^+^ current were also altered with a depolarizing shift of 4 mV in steady-state activation. The RMP of atrial myocytes from *Trpm4*^-/-^ mice was depolarized by 1 mV; however, we did not observe any alteration in I_K1_ in atrial myocytes from *Trpm4*^-/-^ mice. The combined effects of reduced peak Na^+^ currents, altered biophysical properties, and depolarized RMP in *Trpm4*^-/-^ could reduce the total availability of Na_V_1.5 for activation.

Na_V_1.5-mediated Na^+^ current plays a critical role in excitability and conduction velocity in cardiac tissue [36,37]. The strong functional reduction in Na_V_1.5 expression in cardiac myocytes observed in our study was expected to reflect on the whole-heart electrical activity by slowing atrial and ventricular depolarization [38]. Indeed, in our pseudo-ECG recordings on explanted hearts, P duration was longer, and QRS duration shows a trend toward broadening in *Trpm4*^-/-^ hearts. Challenging the explanted hearts with the Na^+^ channel blocker mexiletine confirmed the broadening of QRS in *Trpm4*^-/-^ hearts, which were more sensitive towards mexiletine than WT hearts. In addition to slower depolarization, *Trpm4*^-/-^ hearts also presented significant slowing of intraventricular conduction in the right ventricles compared to WT as observed in intracardiac electrocardiograms. These cellular and whole-heart electrophysiology findings are consistent with a functional reduction of Na_V_1.5 in *Trpm4*-deficient hearts.

In addition, surface ECGs from anesthetized *Trpm4*^-/-^ mice showed a lower heart rate than those from WT mice. Although a direct decrease in heart rate was not observed in the previous studies, the slowing of electrical propagation and sinus pauses were more evident in *Trpm4-’-* mice than in WT [17]. Moreover, in this study, the mice were freely moving and the baseline heart rate was ~500 bpm, while in our case under anesthesia, the heart rate was ~300 bpm. This somehow unmasked the effect of *Trpm4* deletion on the heart rate. A previous study has shown that, in mouse sinoatrial node, Trpm4 activates at lower heart rates, thus acting as an accelerator and increasing the heart rate to counteract bradycardia [39]. However, we did not observe any heart rate change in spontaneously beating *Trpm4*^-/-^ hearts in pseudo-ECGs, where the basal heart rate is even lower (~220 bpm) (Table 2). The effect on heart rate in *Trpm4*^-/-^ mouse surface ECGs is either due to anesthesia or a functional consequence of *Trpm4* deletion in the sino-atrial node, which needs to be cautiously explored in future studies.

Although cardiac phenotypes in *Trpm4^-/-^* mice have been characterized earlier by different groups, none of these studies observed any alteration in peak Na^+^ currents or Na_V_1.5 protein expression [16,17]. This discrepancy between our and other studies could be partially explained by different study designs (isolated myocytes vs. whole tissues, perforated-patch clamp vs. microelectrode / whole-cell recordings). A recent review [40] highlighted that the difference in cardiac phenotype observed in *Trpm4*^-/-^ mice could be strain-dependent. In this study, we showed a similar reduction of peak Na^+^ current due to *Trpm4* deletion in ventricular myocytes from two different mouse lines (Figure 3 and Supplementary 5). Furthermore, we validated the reduction of peak Na^+^ current in whole heart using pseudo-ECGs on mexiletine-perfused explanted hearts and protein expression studies.

To the best of our knowledge, this study provides the first evidence of a functional interaction between Trpm4 and Na_V_1.5 in cardiac tissue. Na_V_1.5 contributes to the fast upstroke in the depolarization phase of a cardiac action potential, while Trpm4-mediated current has been found in the repolarization phase [16–19]. It is intriguing how seemingly distinct depolarizing and repolarizing current components can interact. Yet, a similar concept of ion channel co-regulation has been observed recently for Na_V_1.5 and K_ir_2.1 [41–43]. The authors showed Na_V_1.5, and K_ir_2.1 modulate each other’s function and expression within a macromolecular complex through their respective PDZ-binding domains to regulate cardiac excitability. Another study shows the interaction between hERG and Na_V_1.5 even at the level of mRNA transcripts [44]. The authors demonstrate an interaction of the transcripts of hERG1a, hERG1b, and Na_V_1.5, which regulate each other’s expression at the membrane. Although our study has not addressed whether Trpm4 and Na_V_1.5 directly or indirectly interact at mRNA or protein level, we observed a change in the protein expression and function of Na_V_1.5 due to *Trpm4* deletion without any alteration in *Scn5a* mRNA expression, suggesting a possible interaction and regulation at the post-translational and membrane trafficking levels. This supports our hypothesis that T rpm4 and Na_V_1.5 are interacting protein partners and important for stabilization at the membrane. Nevertheless, future studies to discern possible mechanisms and to determine the level and other partners in the Trpm4-Na_V_1.5 interactions are warranted.

The importance of TRPM4 in cardiac electrical activity is highlighted by mutations in its gene linked to conduction disorders such as right bundle branch block, Brugada syndrome, and atrio-ventricular block[24,40]. Although the first mutation reported in human *TRPM4* was a gain-of-function mutation linked to progressive heart block, several later studies reported both gain- and loss-of-function mutations in *TRPM4* related to different cardiac conduction disorders[11–13,45]. Until now, the mechanisms underlying the genotype-phenotype correlation in TRPM4 mutations remains unclear. Our study suggests a molecular interaction between TRPM4 and Na_V_1.5, brings in a newer perspective on the possible implications of different *TRPM4* mutations found in humans with cardiac conduction disorders. We propose that alterations in TRPM4 expression due to mutations could critically affect the availability of Na_V_1.5 at the membrane. Thus, an increase or decrease in TRPM4 expression may eventually lead to conduction delays, which are indeed observed in *TRPM4* mutation carriers.

In summary, we observed a decrease in Na_V_1.5 expression and function in *Trpm4*-deficient mouse hearts. While the molecular mechanisms underlying this observation are not yet fully understood, one can speculate that the observed overlapping clinical phenotype found in patients with mutations in either *SCN5A* or *TRPM4* may be explained by the co-reg-ulation of Na_V_1.5 expression by the channel TRPM4. Future research addressing this intriguing hypothesis is required.

## Acknowledgments

This work was supported by the Swiss Heart Foundation Grant (grant no 310030_184783) and NCCR TransCure (51NF40-185544 to HA and NCCR Transcure Young Scientist award to OLC). We thank Dr. Rudi Vennekens from the Laboratory of Ion Channel Research, KU Leuven, Belgium, for providing *Trpm4*^-/-^ mice and for providing suggestions and feedback on the manuscript. We thank Dr. Thomas Jespersen, Dr. Morten Bækgaard Thomsen, Dr. Morten Schank Nielsen and Dr. Stefan Sattler from the Department of Biomedical Sciences, University of Copenhagen, Denmark, for providing valuable inputs for intracardiac and pseudo-ECG data analyses. We appreciate Sabrina Guichard from HA group for technical assistance with mouse breeding and genotyping. We acknowledge Dr. Sarah Vermij for proofreading and editing.

## SUPPLEMENTARY MATERIAL

**Supplementary Figure 1:**
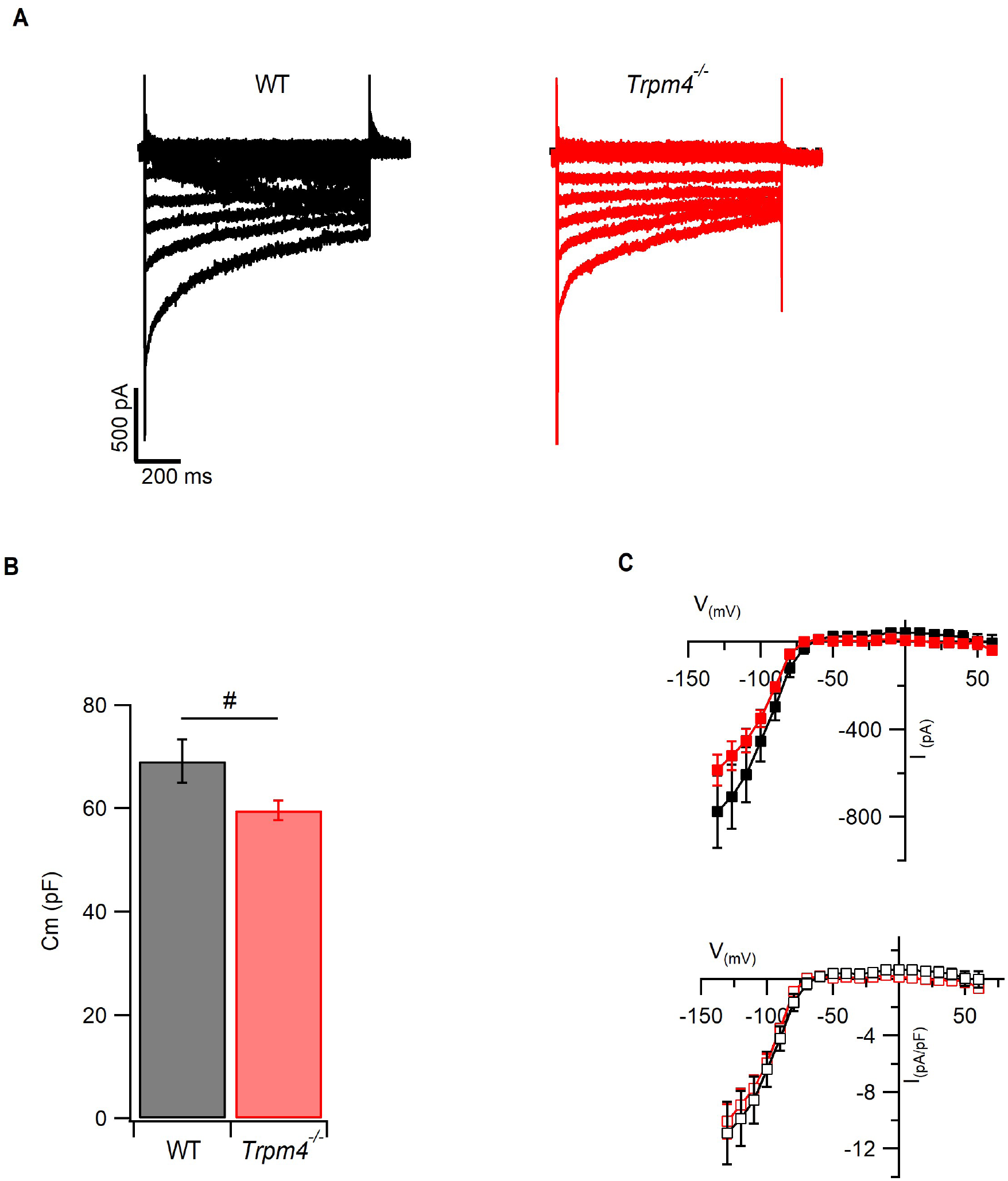
I_K1_ current measurement in atrial myocytes. A) Representative voltage-clamp current traces for assessment of Ba^+^ sensitive inward-rectifier K^+^ current I_K1_ in WT or *Trpm4^-/-^* atrial cardiomyocytes. B) Average cell capacitance measured and compared between both genotypes. C) Current-voltage relationship of peak I_K1_ represented either as current or current density normalized to cell capacitance compared between WT (*N* = 3, n = 18) and *Trpm4^-/-^* (*N* = 3, *n* = 19) mice.

**Supplementary Figure 2:**
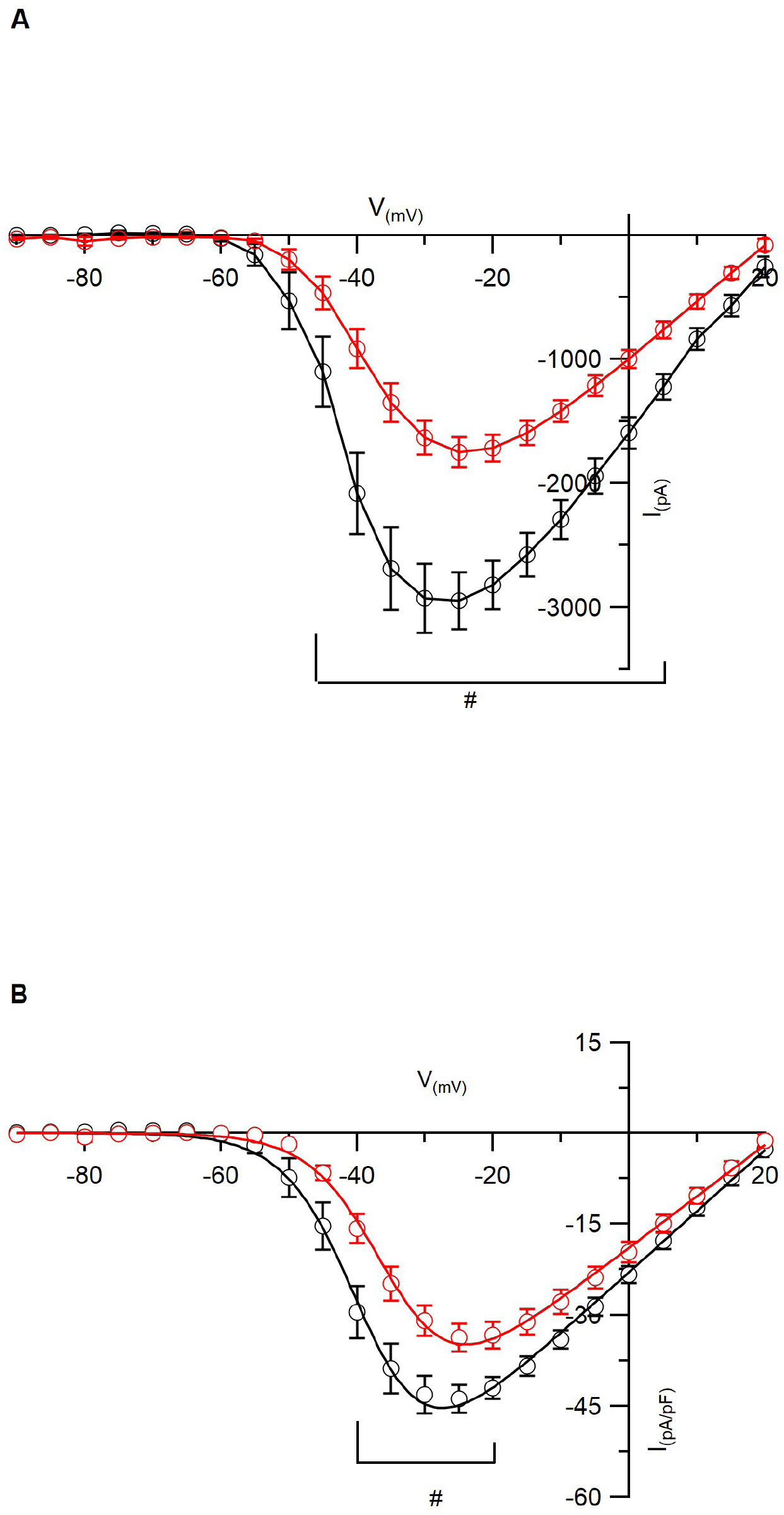
Na_V_1.5 current (A) and current density normalized to cell capacitance (B) of WT and *Trpm4^-/-^* mice (*N* = 5, *n* = 20 WT, *n* = 22 *Trpm4^-/-^*; #: *p* < 0.05, WT vs *Trpm4^-/-^* at a given voltage).

**Supplementary Figure 3:**
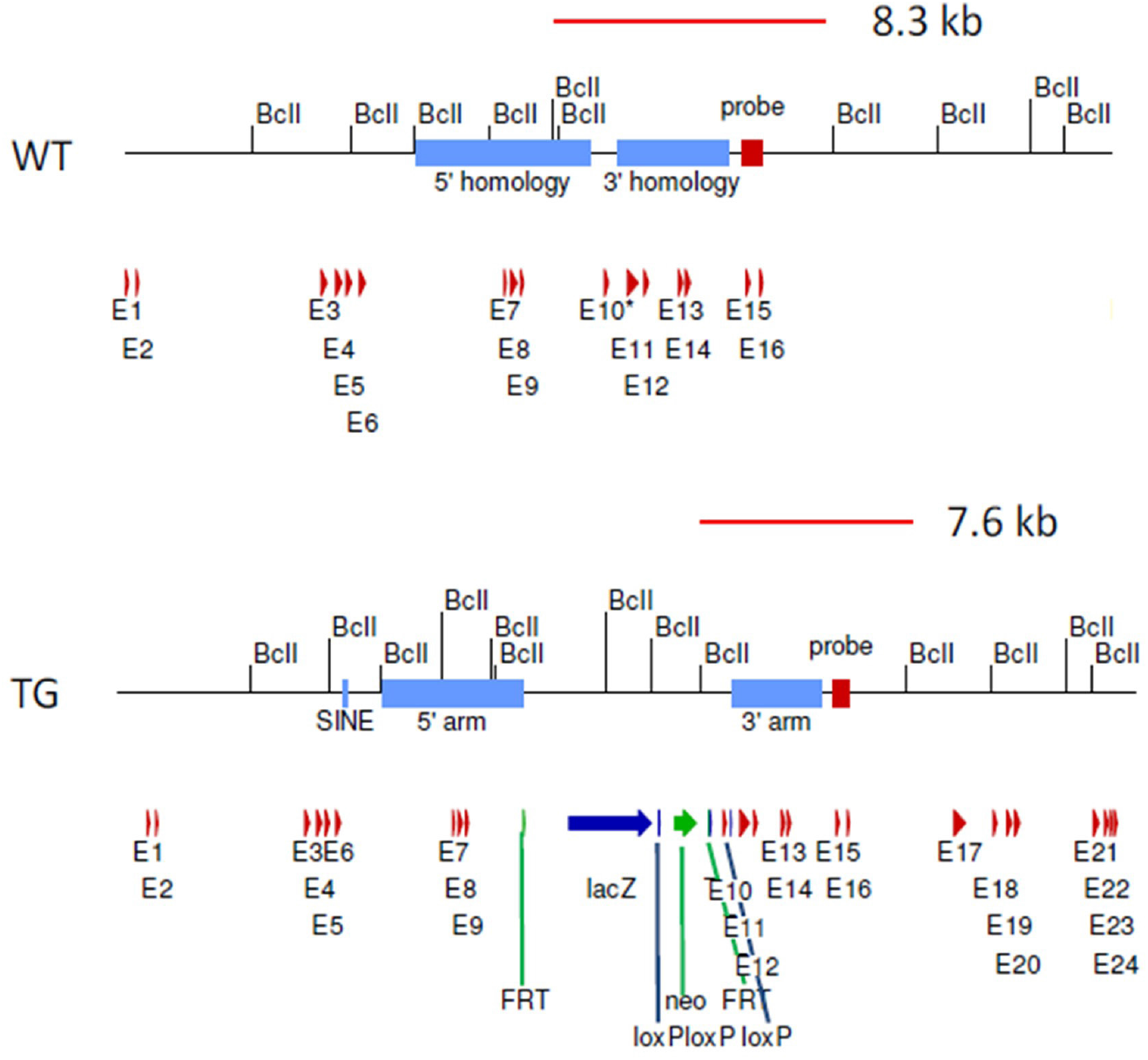
Generation of *Trpm4* knock-out mouse strain. Schematic drawing of the targeted *Trpm4* locus. Two alleles of the *Trpm4* locus are depicted: the wild type locus (WT) and the targeted allele (TG). After homologous recombination, three loxP sites (blue arrows) are inserted into the genome flanking exon 10 of Trpm4 (red arrows). Additionally, an FRT flanked neomycin (green) / lacZ (blue) cassette is inserted upstream of exon 10. NB: this *Trpm4*^-/-^ strain (generated by PolyGene AG, Rumlang, Switzerland) is backcrossed on a C57BL6/J background. In all experiments, male *Trpm4*^-/-^ mice and wildtype (WT) littermates aged 12-15 weeks were used.

**Supplementary Figure 4:**
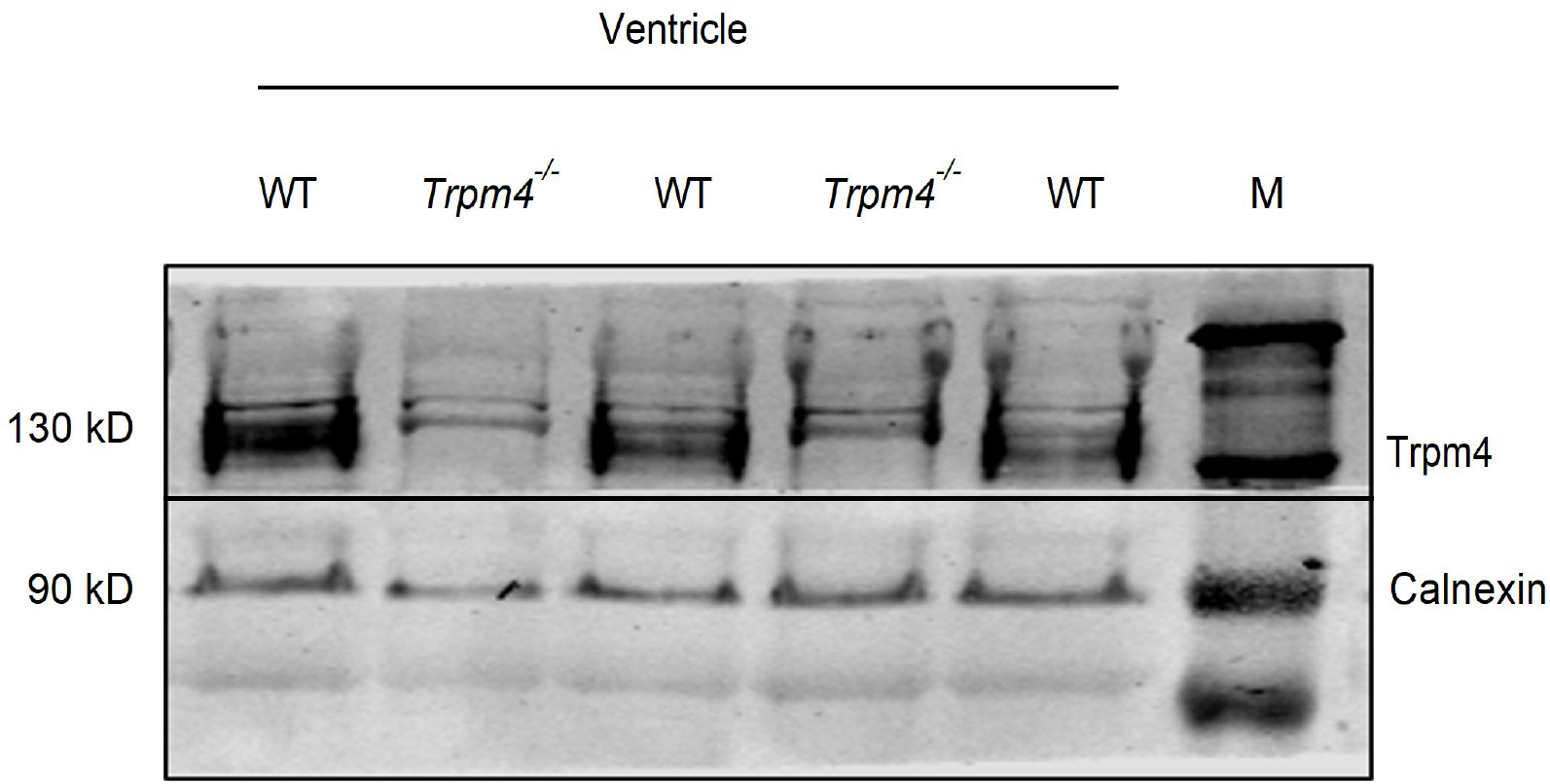
Expression of *Trpm4* in mouse ventricle. Representative immunoblots of Trpm4 total protein expression in mouse ventricular tissue.

**Supplementary Figure 5:**
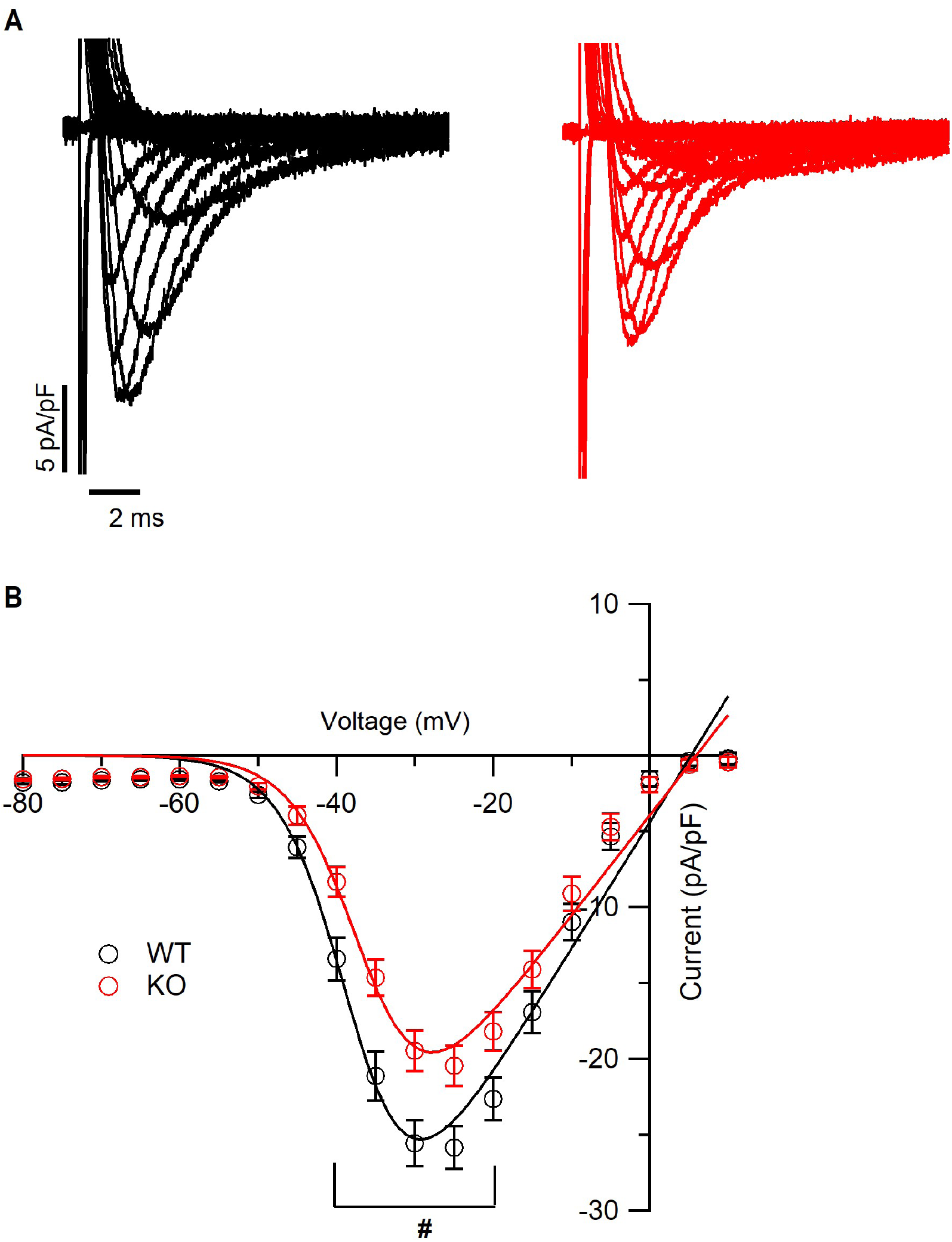
Na^+^ current recordings from isolated cardiomyocytes. A) Representative voltage-clamp current traces for peak Na^+^ current measurements in WT and *Trpm4^-/-^* isolated ventricular myocytes. B) Current-voltage relationship of peak Na^+^ current represented as current density in WT (*N* = 6, *n* = 43) and *Trpm4*^-/-^ (*N* = 6, *n* = 38) mice. (*: *p* < 0.05 at given voltages for WT vs. *Trpm4*^-/-^).

